# Cysteine reactivity profiling identifies host regulators of *Mycobacterium tuberculosis* replication in human macrophages

**DOI:** 10.1101/2025.08.30.673236

**Authors:** John Neff, Kristen E. DeMeester, Paola K. Parraga, Radu Suciu, Melissa Dix, Gabriel Simon, Max A. Gianakopoulos, Bruno Melillo, Benjamin F. Cravatt, Michael U. Shiloh

## Abstract

Innate immune cells such as monocytes and macrophages provide the earliest defense against infection by intracellular pathogens by initiating signaling pathways and restricting pathogen replication. However, the full complement of proteins that mediate cell-autonomous immunity remains incompletely defined. Here, we applied cysteine-directed activity-based protein profiling (ABPP) to map proteome-wide cysteine reactivity changes in THP-1 monocytes and primary human monocyte-derived macrophages during *Mycobacterium tuberculosis* (Mtb) infection. Across both cell types, we quantified 148 cysteine residues with altered reactivity. Genetic perturbation of a subset of proteins harboring these changes significantly impacted Mtb replication, revealing functional links between site-specific cysteine reactivity and antimicrobial defense. These data define previously unrecognized host protein changes during Mtb infection and provide a resource for investigating post-translational events that regulate innate immune responses to intracellular bacteria.

## Introduction

*Mycobacterium tuberculosis* (Mtb), the causative agent of tuberculosis, remains one of the world’s most lethal pathogens, causing 1–2 million deaths annually ^1, 2^. Mtb survives and replicates within host macrophages, and systematic approaches have been used to interrogate this host–pathogen interaction, including global analyses of host transcription, non-coding RNA activity, protein abundance, and post-translational modifications ^3–7^. These studies have identified numerous host defense pathways, yet the role of many post-translational protein changes, particularly those beyond phosphorylation, remains underexplored in the context of Mtb infection.

Activity-based protein profiling (ABPP) is a chemical proteomic strategy that uses reactive probes to covalently bind specific amino acid side chains in native biological systems ^8–10^. This enables the simultaneous detection, identification, and quantification of reactive amino acids and proteins, revealing changes in protein activity, structure, or interactions in response to cellular perturbations ^10^. Cysteine-directed ABPP, using electrophilic probes such as iodoacetamide (IA) linked to an affinity tag, has been widely applied to map reactive cysteines and their ligandability in immune-relevant proteins, including those from human T cells ^11^.

Cysteine-reactivity changes mapped in ABPP experiments can identify peptide-specific and general protein abundance changes (where all cysteine-containing peptides from a given protein coordinately change) ^11^. Site-specific cysteine reactivity changes can help identify post-translational events that regulate protein structure and function, including chemical modifications, conformational shifts, and/or altered binding partner interactions^8^. ABPP can also evaluate how other small molecules react with cysteine residues through competitive blockade of probe engagement^11^. Among various types of electrophilic small molecules studied by competitive ABPP^8^, small molecular weight fragments can provide a broad survey of sites of ligandability in native biological systems ^11–14^.

We hypothesized that cysteine-directed ABPP could be leveraged to map macrophage protein reactivity changes during infection with an intracellular bacterial pathogen and identify novel regulators of antimicrobial defense. To test this, we infected THP-1-derived macrophages ^15^ and primary CD14⁺ monocyte-derived macrophages from healthy donors with Mtb, generating a proteome-wide cysteine reactivity map in each cell type. We then used shRNA-mediated knockdown in THP-1 cells to test the roles of proteins with previously uncharacterized cysteine reactivity changes in the context of human cell-autonomous immunity to Mtb. Our results reveal changes in cysteine reactivity in proteins with known conformational or interactome alterations during Mtb infection. Additionally, several cysteine reactivity changes were found in proteins not previously associated with Mtb pathogenesis that here we demonstrate impact Mtb replication in human macrophages.

## Results

### Global mapping of cysteine reactivity in Mtb-infected human macrophages

We applied cysteine-directed ABPP to profile proteome-wide changes in cysteine reactivity during *Mycobacterium tuberculosis* (Mtb) infection of human macrophages. Two models were used in this platform: PMA-differentiated THP-1 monocytes (hereafter called THP-1 cells) and primary CD14⁺ monocyte-derived macrophages (hereafter called PBMCs). We infected cells with Mtb (MOI 5) for 1 h, then incubated them for either 4 h or 16 h before collecting proteomes for cysteine-directed ABPP using an iodoacetamide-desthiobiotin (IA-DTB) probe^11, 16^ (Fig. 1A). These time points were selected to capture both short and long-term host-cell proteome changes upon infection. The most significant reactivity changes were observed at 16 h (Tables S7–S10) and therefore we focused subsequent analyses on this time point.

**Figure 1:**
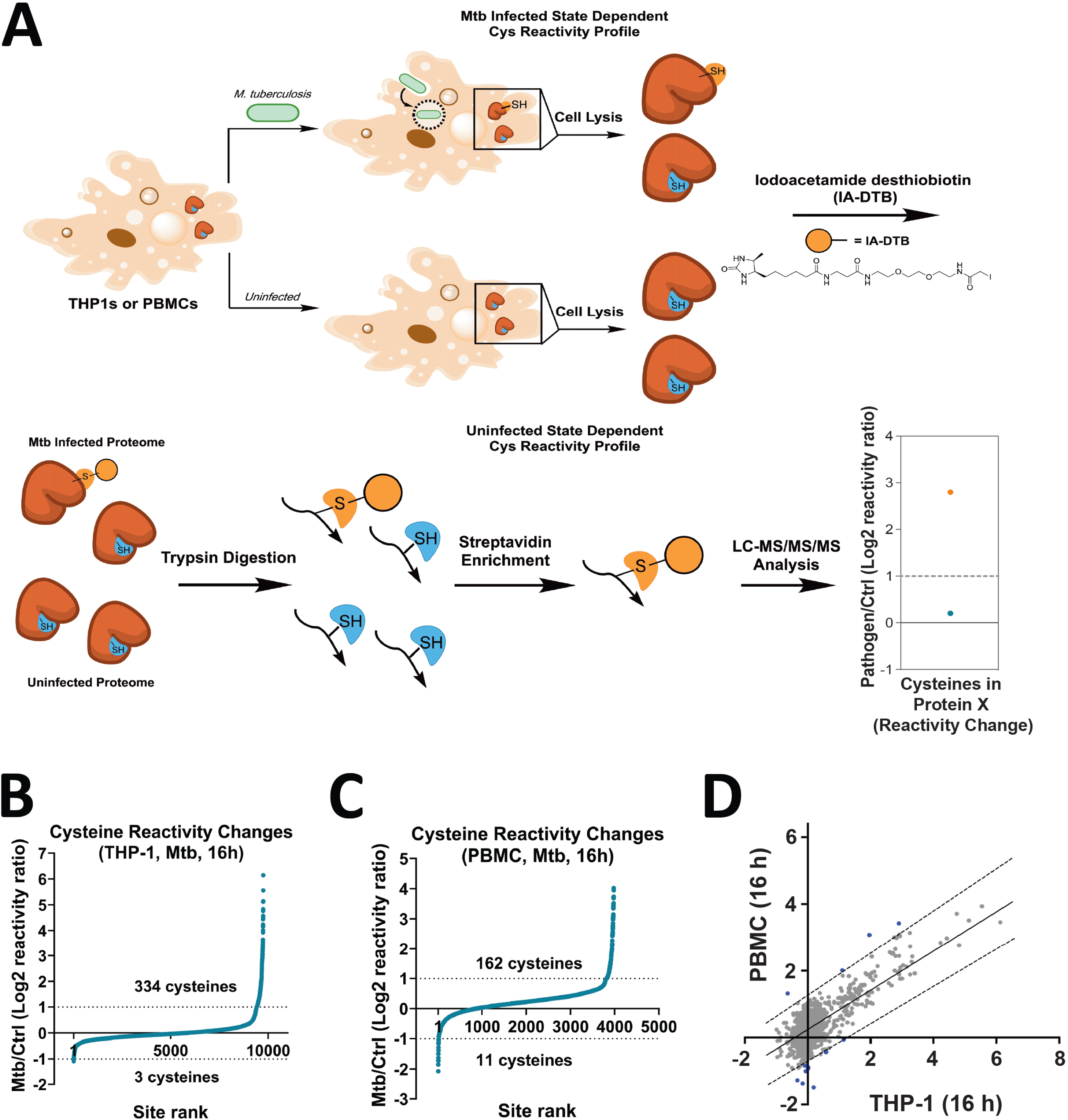
Cysteine-directed ABPP of Mtb-infected THP-1 cells and PBMCs. (A) Graphical representation of the ABPP workflow demonstrating IA-DTB probe binding and enrichment and chemical structure of the IA-DTB probe. . (B, C) Waterfall plots of all cysteines quantified in the ABPP screens in Mtb-infected THP-1 and PBMCs, respectively. Sites shown on the plot are quantified in >= 2 ABPP replicates and if log2R >= 1 then CV<=40% OR if log2R <1, then standard deviation (sd) is <=35. See Table 2 in Supplementary Information. (D) Comparison of proteome-wide cysteine-site reactivities (log2R) in THP-1 cells and PBMCs. Plain line: line of best fit. Dotted lines: 1-log higher than line of best fit (high end of 95% CI) or 1-log lower than line of best fit (low end of 95% CI). Outlier cysteine sites (beyond dotted lines) are shown in blue.

In three independent replicates of 16 h Mtb-infected THP-1 cells, we quantified 18,380 cysteine sites from 6,055 proteins. Of these cysteines, 337 showed either a 50 % increase or decrease in reactivity between uninfected and infected macrophages (Fig. 1B, Table S8). In PBMCs differentiated from five donors, we quantified 13,333 cysteines from 4,927 proteins. Of these cysteines, 173 showed either a 50 % increase or decrease in reactivity between uninfected and infected cells (Fig. 1C, Table S10). The smaller number of altered sites in PBMCs likely reflected donor-to-donor variability, yet the number of unique proteins and reactive sites was comparable to previous ABPP studies with non-infectious perturbations ^11^. To further evaluate the cysteine-reactive proteome, we performed Reactome functional clustering analysis ^17^,and, as expected, proteins involved in response to intracellular pathogens were highly enriched (Tables S11, S12).

We observed broadly similar cysteine reactivity changes between THP-1 cells and PBMCs at 16 h (Fig. 1D). We defined cell model–specific changes as those deviating ≥2-fold from the line of best fit (Fig. 1D, blue). Shared pathway changes included major hallmarks of the macrophage immune response to Mtb infection, including type I interferon signaling ^18, 19^, the OAS antiviral response ^20, 21^, interferon-γ signaling ^4, 22–26^, and NF-kB activity ^27, 28^ (Tables S11,S12). Proteins unique to either model did not cluster into discrete immune pathways, suggesting that differences between THP-1 cells and PBMCs are dispersed across the proteome rather than concentrated in specific processes.

We next analyzed global changes in our cysteine-directed ABPP data to identify proteins showing cysteine reactivity changes without simultaneous alterations in abundance. Proteins with a median change in reactivity that was two-fold or greater were interpreted as changing in abundance (Supplementary Data, Table S5). In Mtb-infected THP-1 cells and PBMCs, 54 and 22 proteins, respectively, showed changes in abundance. Cysteines with reactivity that, upon infection, varied 1.5-fold or more than the median of all sites in the protein were flagged as reactivity rather than abundance changes (Supplementary Data, Table S6. In Mtb-infected THP-1 cells, 25 cysteines from 24 proteins exhibited changes in reactivity, and seven of those cysteines were found in six proteins also changing in abundance (Supplementary Data, Tables S5, S6). In Mtb-infected PBMCs, 116 cysteines from 109 proteins changed in reactivity, and two of those sites were found in two proteins also changing in abundance (Supplementary Data, Tables S5, S6). We identified reactivity changes in two categories of host proteins: those with established roles in innate immunity and those with limited information about their roles in Mtb pathogenesis.

### Cysteine reactivity changes in proteins with established immune roles

We next examined proteins with known functions in innate immunity. In PBMCs, we found that ERAP1, an aminopeptidase critical for trimming peptides for MHC class I presentation ^29^, showed an eight-fold decrease in reactivity of C498 after Mtb infection (Fig. 2A, 2C). As C498 is located within ERAP1’s catalytic domain, this observation suggested reduced catalytic-site accessibility in response to Mtb infection. We also found that NFKB2, the precursor for several components of the NF-κB signaling pathway known to have an important role in Mtb pathogenesis ^28^, increased nearly four-fold in abundance in response to Mtb infection but showed a modest (<2-fold) increase in reactivity of C57 (Fig. 2B, 2D). This observation suggests that C57 may be obstructed in its reactivity with the IA-DTB probe in Mtb infection. After assessing cysteine reactivity changes in known innate immune proteins, we turned our attention to assessing cysteine reactivity changes in proteins with limited roles in Mtb pathogenesis.

**Figure 2:**
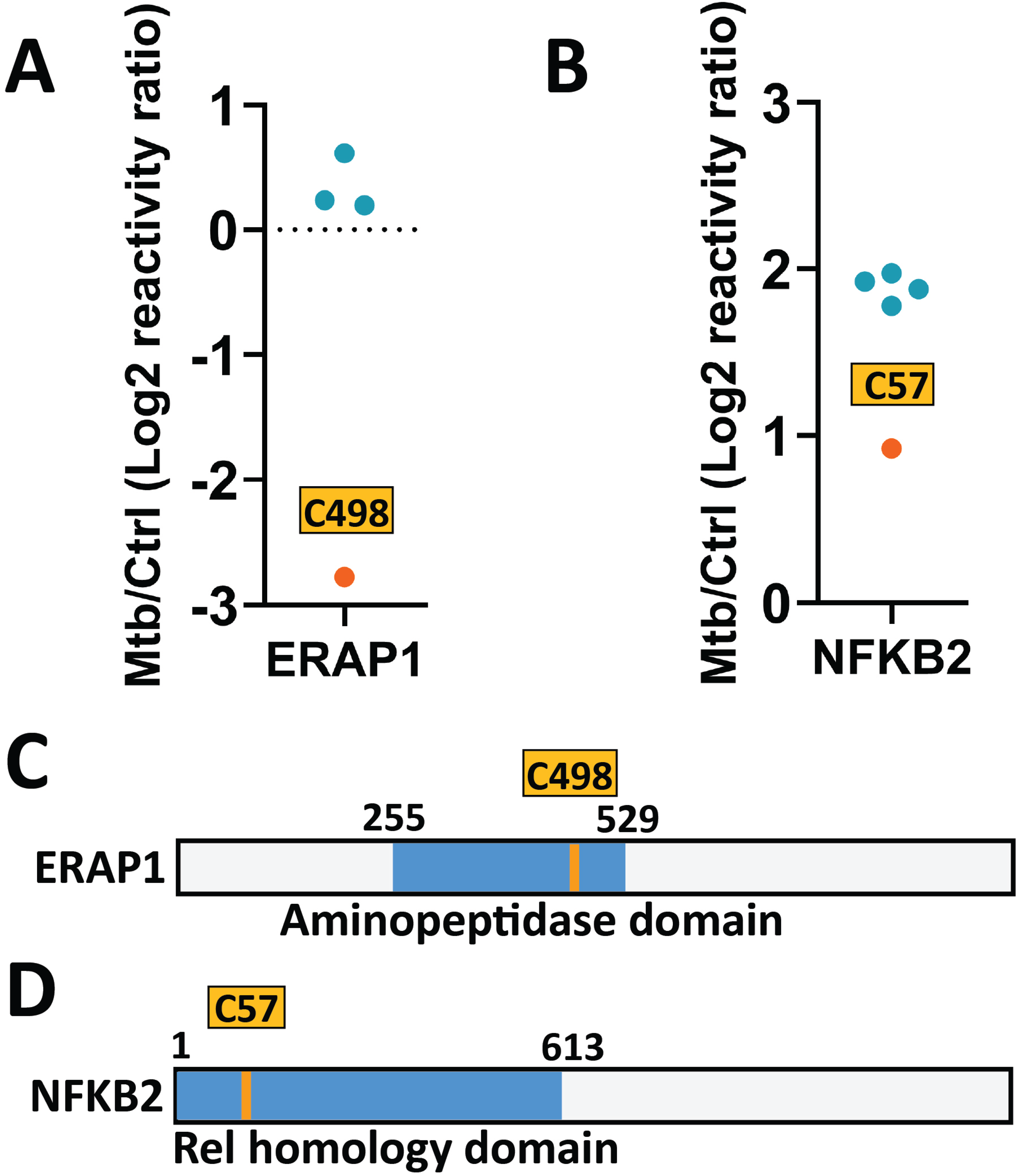
ABPP identifies known protein conformational changes. (A, B) Plot of quantified cysteines of ERAP1 and NFKB2 in Mtb infected macrophages expressed as Log2 reactivity ratio of Mtb/uninfected cells, respectively. (C,D) Graphic cysteine maps of ERAP1 and NFKB2 with relevant domain highlighted in blue and reactive cysteine highlighted in orange

### Novel reactivity changes in lysosome–autophagosome tethering proteins

Regulation of lysosome fusion with Mtb-containing organelles plays a vital role in Mtb pathogenesis^30–32^. We identified novel reactivity changes in two components of the HOPS tethering complex, which mediates lysosome–autophagosome fusion in conjunction with Rab GTPases ^33, 34^. While the HOPS complex has a known role in autophagy, its contribution to Mtb infection remains undefined. We found that two members of the HOPS complex, VPS39 and VPS18, contained cysteines changing in reactivity in response to Mtb infection (Fig. 3A). In VPS39, we observed a >50% increase in reactivity of C852/C855 within a C-terminal zinc-finger domain, without changes in protein abundance in response to Mtb infection (Fig. 3A, 3B). This implies that the zinc-finger domain has increased cytosolic access in response to Mtb infection.

**Figure 3:**
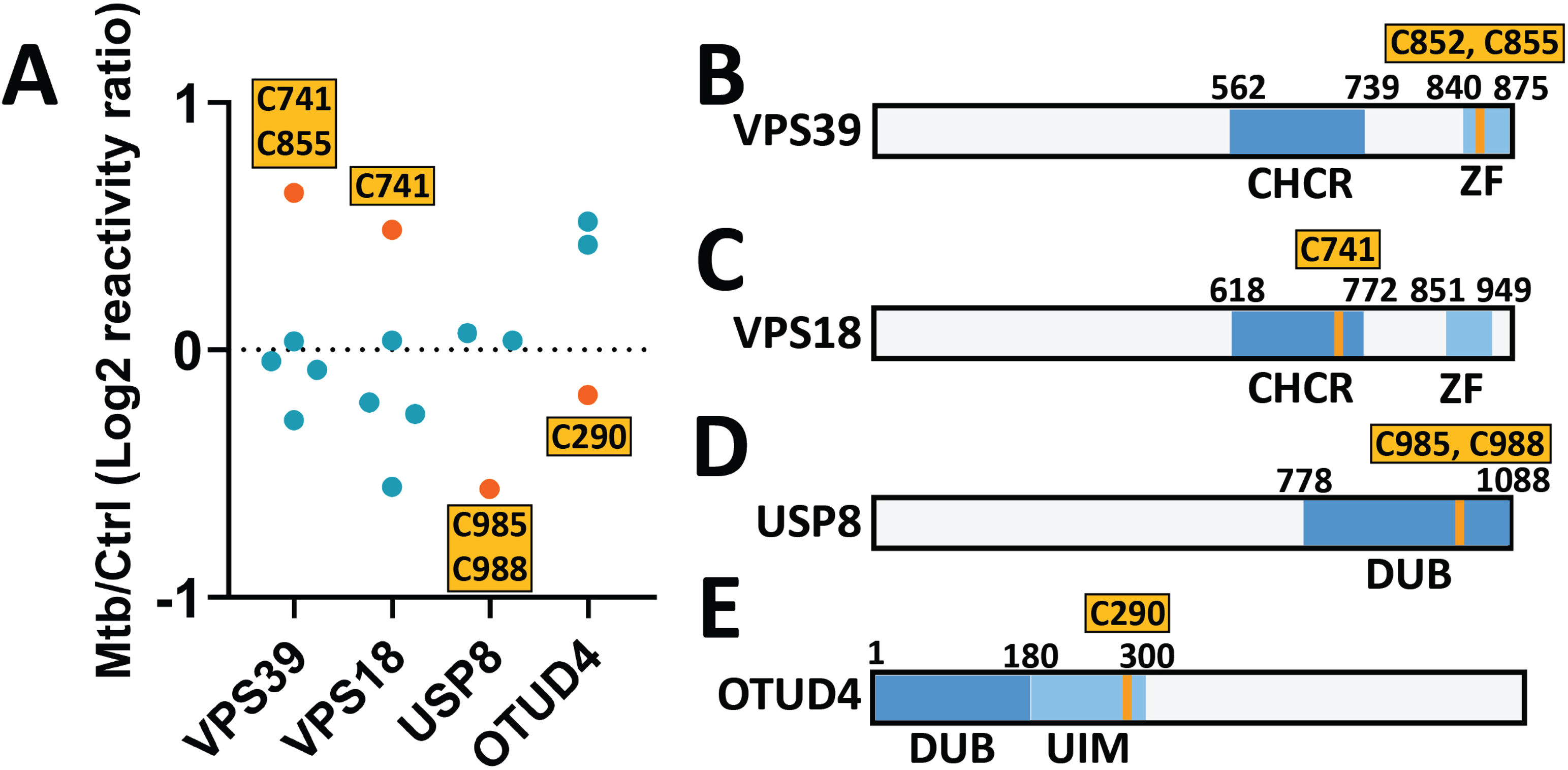
Additional examples of cysteine-reactivity changes in Mtb-infected cells. (A) Plot of quantified cysteines of VPS39, VPS18, USP8, and OTDU4 in Mtb infected macrophages expressed as Log2 reactivity ratio of Mtb/uninfected cells. Graphic cysteine map of VPS39, VPS18, USP8, and OTDU4, respectively with relevant domains highlighted in blue and reactive cysteines highlighted in orange

In VPS18, we detected a 40% increase in reactivity of C741 in response to Mtb infection. VPS18 C742 is within a clathrin heavy chain repeat domain common to multiple members of the HOPS complex that plays an important role in the localization and distribution of the HOPS complex ^35^ (Fig. 3A, C). These changes suggest altered domain accessibility during infection.

### Reactivity changes in deubiquitinating enzymes

We also found infection-associated changes in deubiquitinating enzymes (DUBs), which are less studied in Mtb pathogenesis than ubiquitin ligases and adaptors ^36–39^. In USP8, we detected a 33% decrease in reactivity of C985/C988 within its catalytic domain (Fig. 3A, 3D) in response to Mtb infection as compared to uninfected control. USP8 activity is linked to membrane repair, oxidative stress responses, and inhibition of autophagy during Mtb infection ^40^. In OTUD4, C290 reactivity decreased by 20%, in contrast to 30-40% increases observed at other cysteines within the protein upon Mtb infection (Fig. 3A, 3E). This site lies adjacent to the catalytic domain in a region of the protein that controls ubiquitin linkage specificity ^41^, suggesting a potential infection-induced shift in substrate preference.

### Phenotypic screening identifies cysteine-reactive proteins modulating Mtb growth

To determine whether proteins with altered cysteine reactivity influence Mtb growth, we performed an shRNA screen in THP-1 cells targeting selected proteins from our ABPP datasets. We chose candidates based on both our reactivity mapping and literature searches for potential links to host–pathogen interactions. Each gene was targeted with three to four independent shRNAs (Table S18), and we infected the resulting THP-1 knockdown library with luminescent Mtb (Mtb-lux) to monitor bacterial growth over time (Fig. 4A). We plotted the relative growth of Mtb-lux for each gene in the THP-1 shRNA library using a heat map (Fig. 4B).

**Figure 4:**
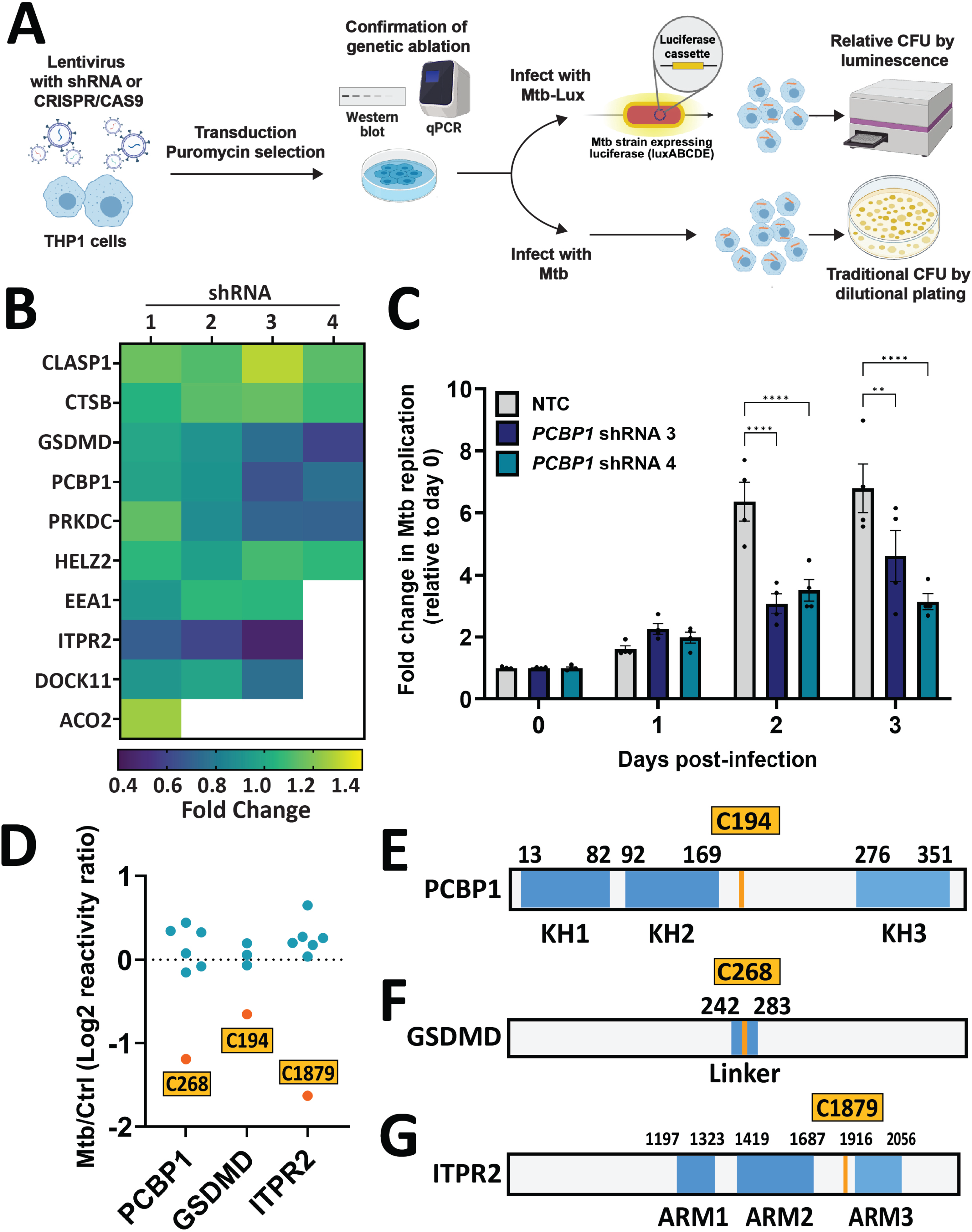
Functional analysis in THP-1 cells of genes whose encoded proteins demonstrate cysteine reactivity changes. (A) Schematic of THP-1 shRNA knockdown mini-library generation and infection with Mtb-lux or Mtb. (B) Heat map of bacterial replication in shRNA knockdown THP-1 lines 5 days post infection with values measured in fold-change with respect to bacterial replication in WT THP-1 cells (C) Time course CFU obtained from *PCBP1* shRNA THP-1 lines infected at an MOI of 1 expressed as fold-change relative to Day 0 (D) Plot of quantified cysteines of GSDMD, PCBP1 and ITPR2 in Mtb infected macrophages expressed as Log2 reactivity ratio of Mtb/uninfected cells. (E-G) Graphic map of GSDMD (D), PCBP1(E) and ITPR2 (F), respectively, with relevant domains highlighted in blue and reactive cysteines highlighted in orange

Knockdown of several genes had a marked effect on bacterial burden. Depletion of *GSDMD* (gasdermin D, a pore-forming gasdermin that induces pyroptosis) ^42^, *PCBP1* (poly(RC)-binding protein 1, a poly(rC)-RNA-binding protein implicated in the pathogenesis of several viruses) ^43–45^, *PRKDC* (protein kinase, DNA-activated, catalytic subunit, a DNA-binding protein kinase involved in DNA damage sensing) ^46^, and *ITPR2* (an inositol 1, 4, 5-trisphosphate-gated calcium channel localized to the endoplasmic reticulum) ^47^ reduced Mtb growth compared with control cells, whereas knockdown of *CLASP1* (a regulator of microtubule dynamics) ^48^ enhanced bacterial replication (Fig. 4B). We selected PCBP1, GSDMD, and ITPR2 for further analysis because their knockdown consistently reduced bacterial growth across experiments.

PCBP1 encodes an RNA-binding protein involved in multiple cellular processes, including regulation of innate immunity ^49–51^. Knockdown of *PCBP1* in THP-1 macrophages significantly reduced Mtb growth in both the Mtb-lux assay and conventional CFU measurements (Fig. 4B, 4C). Notably, we detected a >50 % decrease in reactivity of C194 following Mtb infection (Fig. 4D). This residue is located outside of the RNA-binding K-homology domains (Fig. 4E), suggesting that infection may alter PCBP1 function through a site-specific modification distinct from its canonical RNA-binding interface.

GSDMD is a pore-forming protein that mediates pyroptotic cell death upon cleavage by caspases. In Mtb-infected macrophages, C268 reactivity decreased by nearly 50% (Fig. 4D). This residue lies within the linker connecting the active N-terminal and autoinhibitory C-terminal domains and is positioned near both a known protein-protein interaction (PPI) interface ^52^ and Asp275, the caspase-1 cleavage site ^42^ (Fig. 4F). The observed reduction in reactivity could result from caspase-1 mediated cleavage, which occurs during Mtb infection of mouse macrophages ^53^, or altered accessibility of this site. Knockdown of *GSDMD* reduced bacterial replication by approximately 40%, supporting a role for this protein in the intracellular growth environment of Mtb.

ITPR2, a member of the inositol 1,4,5-trisphosphate receptor family, mediates calcium release from the endoplasmic reticulum ^54^ and contributes to calcium-dependent cellular processes, including immune signaling ^55^. We observed a ∼50 % decrease in reactivity of C1879 following Mtb infection (Fig. 4D). This cysteine lies in the middle of three armadillo repeat domains (ARM) (Fig. 4G), structures known to facilitate protein–protein interactions and organelle tethering ^56^. The loss of reactivity may reflect infection-induced recruitment of binding partners ^57, 58^ that occlude IA-DTB probe access, or oxidative modification of this cysteine ^59^, which could alter channel function. Knockdown of *ITPR2* significantly reduced Mtb replication in THP-1 cells, suggesting a role for this calcium channel in supporting bacterial growth.

### Fragment ligandability mapping in infected macrophages

Finally, we used fragment-based ABPP ^11, 13, 14^ to identify ligandable cysteines in Mtb-infected macrophages. During the final hour of infection, we treated cells with the electrophilic “scout fragment” KB05 ^13^ (Fig. 5A), which contains an electrophilic acrylamide warhead for covalent cysteine modification and non-covalent elements to enhance binding. At 16 h post-infection, we identified 74 KB05-liganded cysteines in THP-1 cells and 53 in PBMCs (Tables S13–S17). We included only sites where other cysteines in the same protein showed <25 % IA-DTB blockade to avoid abundance confounding. KB05-liganded cysteines were found in several immune-relevant proteins. One liganded site we identified was C287 of TNF receptor associated factor 2 (TRAF2), a protein involved in NF-kB signal transduction ^60^ (Fig. 5B). Importantly, the 10 amino acid sequence between residues 283-293 of TRAF2 bind directly to Baculoviral IAP Repeat Containing (BIRC) 2 and BIRC3 (Fig. 5D), and this binding is required for the regulation of NF-kB signaling by TRAF2^61, 62^. Another liganded site was C55 of BCL2A1, an anti-apoptotic protein implicated in several forms of cancer ^63, 64^. BCL2A1 C55 is located in a protein region targeted by small molecule inhibitors ^65, 66^ (Fig. 5E). These findings validate the use of fragment-based ligandability mapping in primary human macrophages infected with Mtb and reveal potential targets for covalent probe development.

**Figure 5:**
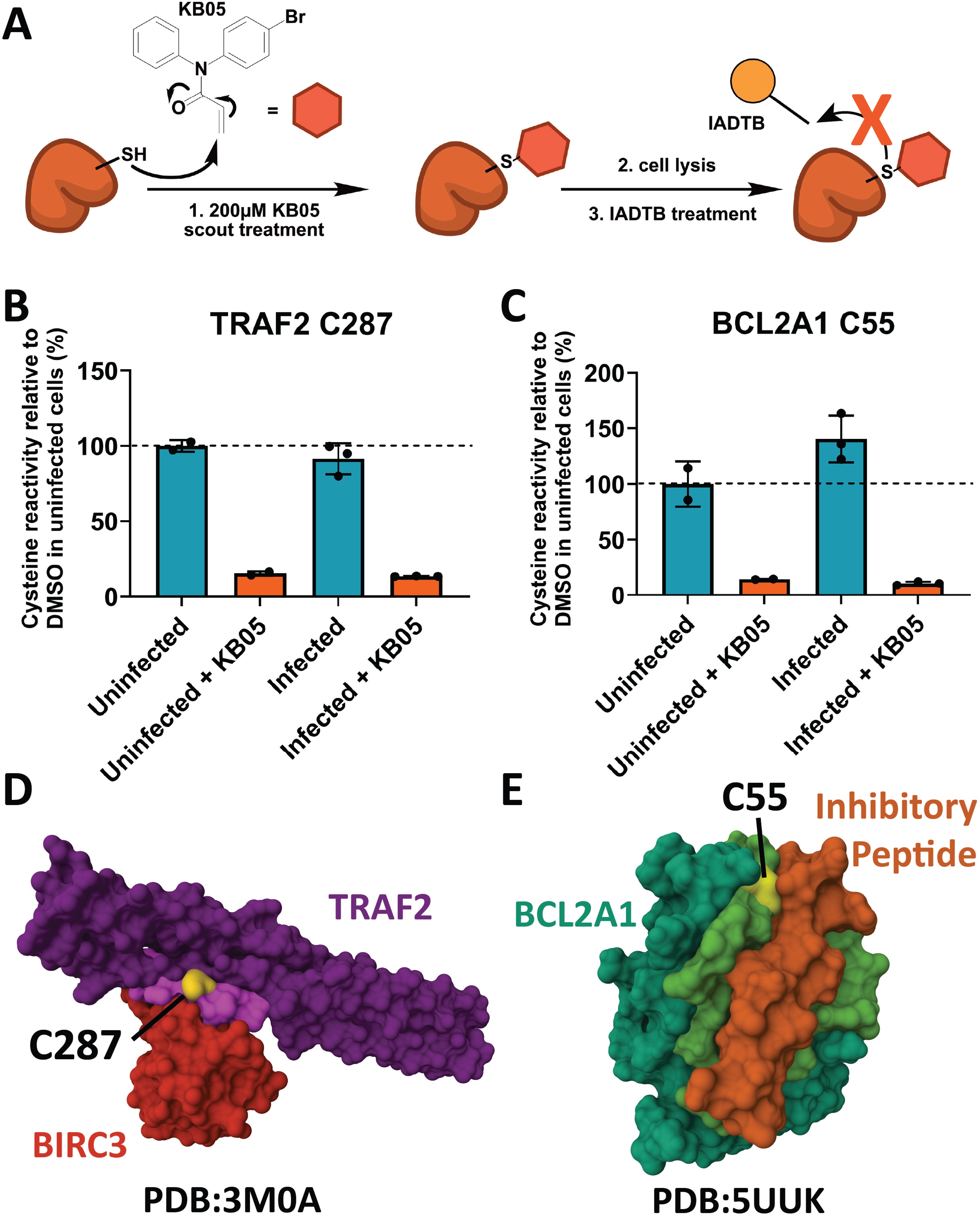
Mapping ligandable cysteines in Mtb-infected cells. (A) Schematic of cysteine protection using a scout fragment (B, C) Bar graphs of MS signal corresponding to percent cysteine reactivity relative to DMSO in uninfected cells of C287 in TRAF2 and C55 of BCL2A1, respectively, with and without pretreatment with KB05 scout fragment. (D) Crystal structure of TRAF2 (purple) bound to BIRC3 (red) with C287 highlighted in yellow. The protein-protein interaction interface of TRAF2 is colored in magenta. All residues of TRAF2 within 5 angstroms of BIRC3 were highlighted as part of the protein-protein interaction interface. (E) Crystal structure of BCL2A1 (blue) in complex with a BCL2A1-specific selected peptide (orange) with C55 highlighted in yellow. The peptide binding groove of BCL2A1 is colored green. All residues of BCL2A1 within 5 angstroms of the depicted inhibitory peptide were highlighted as part of the protein-protein interaction interface.

## Discussion

Post-translational modulation of protein structure and function is a key mechanism by which cells respond to infectious agents. While activity-based protein profiling (ABPP) has been used to detect post-translational changes in amino acid reactivity in multiple systems and cell types^11, 67, 68^, it has not previously been applied to Mtb infection. Here, we provide the first proteome-wide maps of cysteine reactivity in Mtb-infected human macrophages, quantifying more than 15,000 reactive cysteines across over 5,000 unique proteins in both immortalized and primary macrophages. Our analysis captured cysteine reactivity changes in known immune regulators such as ERAP1 and NF-κB, as well as in proteins with less-characterized roles in Mtb infection, including HOPS complex components and deubiquitinases USP8 and OTUD4. We further demonstrated that proteins such as PCBP1, GSDMD, and ITPR2 contribute to Mtb replication in human macrophages. These findings establish cysteine-directed ABPP as a powerful approach for dissecting dynamic protein function during infection.

ABPP offers mechanistic insights that cannot be obtained from transcriptomic or abundance-based proteomics alone ^61^. Many proteins undergo conformational or chemical modifications that alter their activity without changing overall expression levels. ABPP detects such state-dependent changes directly at reactive amino acids, complementing other omics approaches. However, proteins lacking reactive cysteines will not be detected by cysteine-directed ABPP, and certain post-translational changes may be better probed by targeting other residues such as lysine^69, 70^, tyrosine^71^, aspartate/glutamate^72^, methionine^73^, or tryptophan^74^. In our study, we distinguished reactivity shifts from abundance changes by comparing cysteines within the same protein, but confidence decreases when few sites are quantified. Integrating cysteine-directed ABPP with parallel abundance-based proteomics could further strengthen these assignments.

We also applied cysteine-directed ABPP for ligandability mapping using a promiscuous electrophilic scout fragment, revealing numerous cysteines in immune-relevant proteins with potential for covalent targeting. These results provide an entry point for chemical probe or inhibitor development, although conversion of fragment hits into potent, selective compounds will require iterative medicinal chemistry and structural biology. The absence of scout fragment binding does not rule out ligandability by other chemistries, underscoring the need for broader electrophile libraries and orthogonal validation.

Our genetic analysis linked cysteine reactivity changes to functional consequences for Mtb replication. While some knockdowns, such as *CTSB*, did not affect bacterial growth despite prior reports ^75^, these discrepancies may reflect differences in Mtb strain (Erdman vs H37Rv), depletion method (shRNA vs siRNA), or assay readout. Other proteins, including HELZ2 and DOCK11, may regulate infection through immune signaling pathways such as inflammatory signaling and cytokine production, which are not captured by in vitro CFU assays ^76, 77^.

Notably, knockdown of GSDMD reduced Mtb replication by ∼40%. We identified Cys268, near the caspase-1 cleavage site, as a site of decreased reactivity during infection. This location aligns cleavage-dependent activation of GSDMD and the targeting of mitochondria by its N-terminal fragment in murine ^53^ and human ^78^ macrophages, which can drive necroptosis or pyroptosis, respectively. While *Gsdmd* deletion in mice does not alter Mtb burden ^79^, our data suggest that GSDMD negatively regulates cell-autonomous immunity in human macrophages. Such findings point to host proteins whose inhibition, either by small molecules or degraders, could enhance bacterial clearance ^80, 81^.

In summary, we used cysteine-directed ABPP to map Mtb infection-induced changes in the reactive proteome of human macrophages, linking site-specific cysteine reactivity to mechanistic roles in innate immune defense. Our data reveal proteins whose reactivity changes likely reflect altered complex formation, post-translational modification, or structural remodeling during infection. Ligandability maps further identify cysteine sites suitable for covalent probe development, offering a resource for chemical biology approaches to dissect and modulate host–pathogen interactions. Expanding such analyses to diverse electrophilic molecules ^11, 82^ may yield selective covalent ligands that both reveal functional roles of key host proteins and provide starting points for host-directed therapeutic development. By linking infection-induced cysteine reactivity to functional consequences for bacterial replication, we identify new host pathways and ligandable sites that can be exploited to dissect, and potentially reprogram, macrophage immunity against Mtb.

## Materials and Methods

### Bacterial strains and culture

All experiments used the Mtb Erdman strain, which was tested for the presence of PDIM using mass spectrometry. For luminescence experiments measuring Mtb replication, an Mtb Erdman strain expressing the entire luciferase operon (Addgene plasmid #26161) was used (hereafter called Mtb-lux). For macrophage infection experiments, bacteria were grown to an OD600 of 0.5-0.8 at 37°C on a roller apparatus in 7H9 media supplemented with 20% Middlebrook Oleic Albumin Dextrose Catalase Growth Supplement (OADC), 1% glycerol, and 0.1% Tween 80, washed three times with PBS, sonicated at 90% amplitude four times for seven seconds each in a qSonica q125 sonicator with cup horn attachment, and then centrifuged at low speed (300g) to pellet bacterial aggregates. The OD_600_ of the supernatant was then measured and used to calculate the amount of culture needed for the correct MOI assuming that an OD_600_ of 1 is equivalent to a bacterial concentration of 3×10^8^ CFU/mL.

### DNA constructs and lentivirus production

shRNA-expressing lentiviral vectors were purchased from the MISSION shRNA library (Sigma Aldrich). All shRNA sequences are provided in Supplementary Table 18. shRNA knockdown lines were generated using transduction via lentiviruses containing shRNA constructs. Lentiviruses were generated by transfecting HEK293T cells using the Fugene6 transfection reagent with the shRNA construct, the psPAX2 packaging vector (AddGene Plasmid #12260), and the envelope-expressing pMD2.G vector (AddGene Plasmid #12259). Lentivirus was harvested three days post-transfection and added to WT THP-1 cells. After two days of incubation with lentivirus, newly created cell lines were selected for one week using 15ug/mL of blasticidin.

### Mammalian cells and culture

THP-1 cells were cultured in RPMI supplemented with 10% FBS, 1% HEPES, and 1% sodium pyruvate. THP-1 cells were differentiated for two days in 100ng/mL PMA and allowed to rest after a media change to RPMI without PMA for an additional two days before use. Human PBMCs were isolated using Sepmate 50 tubes (StemCell catalog #85450) and the accompanying protocol followed by CD14+ cell enrichment using human CD14 micro beads (Miltenyi catalog #130-050-201) with MACS LS columns and quadroMACS separation magnet (Miltenyi). PBMCs were obtained from buffy coats from anonymous donors provided by a local blood bank. PBMCs were differentiated for 24 hours in RPMI supplemented with 10% FBS, 10% heat inactivated donor-specific serum isolated from blood, 1% HEPES, 1% sodium pyruvate, and 50ng/mL GM-CSF (PeproTech catalog #300-03). Media was then changed to RPMI supplemented with 10% FBS, 1% HEPES, 1% sodium pyruvate, and 50ng/mL GM-CSF and cells were incubated for three days. Finally, the media was changed to RPMI supplemented with 10% FBS, 1% HEPES, and 1% sodium pyruvate and cells were allowed to incubate for three additional days before use in experiments. No antibiotics were included in the media used for macrophage infections.

### ABPP macrophage infection

Bacteria were resuspended in RPMI media with 10% FBS and then added to cells and allowed to infect for one hour at 37°C in a CO_2_ incubator at an MOI of 5. Cells were then washed twice with PBS and fresh media was added. For ABPP experiments, after 15 hours, cells were treated with either RPMI+DMSO or RPMI+200uM KB05 scout fragment. Cells were trypsinized and harvested 1 hour after scout fragment addition and lysed by sonication on ice using a Bioruptor Plus (Diagenode) set to high for five minutes with a 30 second on/off cycle.

### ABPP sample collection and cysteine labeling

After 16 hours, uninfected or infected THP-1 cells or PBMCs were trypsinized for ten minutes at room temperature. The trypsin was then neutralized with an equal volume of RPMI + 10% FBS, cells were removed from the plates using a disposable cell scraper and washed once with 10mL of PBS. Cells were then lysed in 500uL of PBS containing an EDTA-free protease inhibitor cocktail (Sigma #11836170001). Protein content of the samples was quantified using a Pierce BCA Assay kit and samples were diluted to a concentration of 2 mg/mL of total protein in 500uL of PBS plus protease inhibitor. To label reactive cysteines, 5uL of a 100x stock of Iodoacetamide-Desthiobiotin (IA-DTB) was added to each sample to a final concentration of 100uM. Samples were mixed by inversion and incubated for 1 hour at room temperature while protected from light.

To precipitate protein, 600µL MeOH, 100µL CHCl3, and 100µL water were added to the samples. Samples were then vortexed and centrifuged at 16,000xg at 4°C for 15 minutes. All liquid in the samples was aspirated, leaving only a protein disk. To wash, 600µL MeOH and 100µL CHCl3 were added to samples, vortexed, and then centrifuged again under the same conditions. The wash was then aspirated, and pellets were allowed to air dry before being stored at -80°C.

### ABPP sample digestion

90 µL of buffer containing 9M urea, 10 mM DTT and 50 mM triethylammonium bicarbonate buffer (TEAB) was added to each protein pellet sample collected after infection and cysteine labeling. Once the protein disks became translucent and partially dissolved, the tubes were gently mixed by flicking. Samples were heated to 65°C (with shaking after ten minutes by gently flicking the tube as above) for 20 minutes or until samples were completely dissolved. After cooling to room temperature, 10 µL of 500 mM iodoacetamide was added to each sample. Samples were then vortexed and shaken on a rotary shaker at 37 °C for 30 min. 300µL of 50mM TEAB were then added to each sample, followed by 1μg of trypsin (20µg trypsin resuspended in 60µL trypsin buffer and 20µL 100mM CaCl2, with 4µL added to each sample). Samples were then digested overnight at 37°C.

### Biotin-tagged peptide enrichment

300µL of wash buffer (50mM TEAB, 150mM NaCl, 0.2% NP-40) containing 50µL of streptavidin bead slurry (Thermo cat # 20353) was added to digested samples and rotated for two hours at room temperature. Bead slurry was prepared beforehand with 2 washes of 10mL wash buffer each and resuspended at a ratio of 50µL agarose slurry per 300µL of wash buffer. Beads were filtered using a filter spin column (pre-washed with 1ml H_2_O) and were washed 3 times with 1 mL of modified wash buffer (containing only 0.1% NP-40), 3 times with 1 mL PBS, and 3 times with 1 mL with HPLC grade H_2_O. Spin columns were then transferred to low protein-binding eppitubes and gravity eluted by addition of 300 µL of solution containing 50% HPLC grade acetonitrile and 0.1% HPLC grade formic acid into a new low binding tube. This was repeated once for 600uL total elution. A pipette bulb was used to expunge any remaining liquid from the beads, and the eluate was then evaporated to dryness in a speed vac.

### TMT Labeling

Enriched peptides were resuspended in 100μl 200mM EPPS (pH 8) with 30% dry HPLC grade acetonitrile, vortexed, and centrifuged at high speed in a tabletop centrifuge. Samples were then sonicated in a water bath for 5 minutes, vortexed, and centrifuged again. 3ul of the corresponding TMT tag was added to each tube. Samples were then vortexed and left at room temperature for 1 hour. 3μl of 5% hydroxylamine was added to each sample, which were then vortexed and left at room temperature for 15 minutes. After addition of 5μl of formic acid, samples were centrifuged and each 10-plex was combined in a new low binding 2 ml tube. The original tubes were centrifuged again to ensure no sample remained. Combined samples were then evaporated to dryness in a speed vac.

### Desalting using Sep-Pak C18 Cartridge

TMT-labeled samples were resuspended in 500μl of buffer A (95% H20, 5% acetonitrile, 0.1% formic acid) and an additional 20μl of formic acid was added to ensure samples were acidic. Samples were then sonicated in a water bath for 5 minutes. 2. Sep-Pak cartridges were conditioned by adding 1 mL 100% acetonitrile three times and then equilibrated by adding 1 mL of Velos buffer A (95% H20, 5% acetonitrile, 0.1% formic acid) three times. Samples were loaded at a rate of 1 drop/sec and the flow-through was collected and loaded a second time. Samples were then desalted by passing 1 mL of Velos buffer A through the cartridge three times. Samples were eluted by adding 1 mL 80% acetonitrile/0.1% formic acid and cartridge was blown dry. The elution was then dried using a speed vac and resuspended in 500ul of buffer A to load on HPLC for offline high pH fractionation.

### HPLC fractionation

Desalted samples were sonicated in a water bath for 5 minutes, centrifuged at high speed in a tabletop centrifuge for 1 minute, and inspected to ensure no particulate matter was present. The columns were washed in 80% buffer B (80% ACN, 20% water, 0.1% FA) and then equilibrated with 100% buffer A. 96 well 1mL deep plates were prepared for fraction collection by adding 20uL of 20% formic acid. After fraction collection the 96-well plate was dried using a speed vac and resuspended in Velos buffer B, combining each column into a single sample for injection into an Orbitrap Fusion mass spectrometer.

### Mass Spectrometry

The following Mass Spectrometry Peptide Data Collection mirrors Njomen et. al TMT liquid chromatography-mass-spectrometry (LC-MS) analysis^82^. Briefly, samples were analyzed by liquid chromatography tandem mass-spectrometry using an Orbitrap Fusion mass spectrometer (Thermo Scientific) coupled to an UltiMate 3000 Series Rapid Separation LC system and autosampler (Thermo Scientific Dionex), as previously reported^11^. Data was acquired with Thermo Scientific Xcalibur software version 2.2. The peptides were eluted onto a capillary column (75 μm inner diameter fused silica, packed with C18 (Waters, Acquity BEH C18, 1.7 μm, 25 cm)) or an EASY-Spray HPLC column (Thermo ES902, ES903) using an Acclaim PepMap 100 (Thermo 164535) loading column, and separated at a flow rate of 0.25 μL/min. Data was acquired using an MS3-based TMT method on Orbitrap Fusion or Orbitrap Eclipse Tribrid Mass Spectrometers. Briefly, the scan sequence began with an MS1 master scan (Orbitrap analysis, resolution 120,000, 400−1700 m/z, RF lens 60%, automatic gain control [AGC] target 2E5, maximum injection time 50 ms, centroid mode) with dynamic exclusion enabled (repeat count 1, duration 15 s). The top ten precursors were then selected for MS2/MS3 analysis. MS2 analysis consisted of quadrupole isolation (isolation window 0.7) of precursor ion followed by collision-induced dissociation (CID) in the ion trap (AGC 1.8E4, normalized collision energy 35%, maximum injection time 120 ms). Following the acquisition of each MS2 spectrum, synchronous precursor selection (SPS) enabled the selection of up to 10 MS2 fragment ions for MS3 analysis. MS3 precursors were fragmented by HCD and analyzed using the Orbitrap (collision energy 55%, AGC 1.5E5, maximum injection time 120 ms, resolution was 50,000). For MS3 analysis, we used charge state–dependent isolation windows. For charge state z = 2, the MS isolation window was set at 1.2; for z = 3–6, the MS isolation window was set at 0.7. Raw data was acquired and moved to mass spectrometry peptide data processing and analysis.

### Data Processing

The raw MS data files were uploaded to and converted by Integrated Proteomics Pipeline (IP2, v 6.7.1) available at (http://ip2.scripps.edu/ip2/mainMenu.html). MS2 and MS3 files extracted from the raw files using RAW Converter (version 1.1.0.22, available at http://fields.scripps.edu/rawconv/). The data files were then processed using the ProLuCID program based on a reverse concatenated, non-redundant version of the Human UniProt database (release 2016–07). Cysteine residues were searched with a static modification for carboxyamidomethylation (+57.02146 Da). N-termini and lysine residues were searched with a static modification corresponding to the TMT 10 plex tag (+229.16293 Da). To search for the cysteine IA-DTB labeling, a dynamic modification (+398.25292 Da) was used with a maximum number of 2 differential modifications. The census output files from IP2 were further processed by aggregating TMT reporter ion intensities to obtain signals based on unique peptides that are further annotated with protein-cysteine residue numbers. Peptides were required to be at least 6 amino acids long. ProLuCID data was filtered through DTASelect (version 2.0) to achieve a peptide false-positive rate below 1%. The MS3-based peptide quantification was performed with reporter ion mass tolerance set to 50 ppm with Integrated Proteomics Pipeline (IP2). The resulting data were then median normalized per TMT channel and log2 fold changes between the native versus denatured conditions were calculated for each cysteine.

Raw proteomic datasets (.raw) were searched using the Integrated Proteomics Pipeline (IP2 v6.0.2) with Human UniProt database (release 2016–07) (.fasta) as the reference proteome. The IP2 output files (.txt, one file per TMT-plex replicate, see [experiment index in supplement]) comprise lists of raw peptide-spectrum matches (PSMs), each associated with 10 MS3 intensities (i.e., 10 channels) reporting on the abundance of the matched peptide under a given experimental condition. Raw PSMs were considered quantified if control channels (DMSO-treated cells, no pathogen) showed low variability (CV = sd /mean < 0.5) and sufficient intensity (sum signal > 5,000 per channel). Quantified PSMs were further analyzed as outlined in Supplementary Table 2 and described below to identify significant changes in cysteine-site reactivity or protein abundance upon stimulation, as well as ligandable cysteine sites. Pathogen-induced changes in cysteine reactivity and protein abundance. Channel intensities in quantified PSMs were normalized using the control intensities as reference ([DMSO, no pathogen] = 100). Normalized intensities within each channel were then averaged across quantified PSMs associated to the same cysteine site to obtain raw site data. A reactivity ratio R was defined as the ratio of normalized intensity of a given condition to that of control. Lowest-variability quantified sites were those detected in 2 or more ABPP replicates and either (i) log2Rsite ≥ 1 and CV ≤ 0.4, or (ii) log2Rsite < 1 and sd ≤ 35. Channel values for lowest-variability quantified sites were averaged within each experimental condition and grouped by protein. Further analyses required that 3 or more sites were quantified in each protein. If the absolute difference between the reactivity change in one cysteine site and the median reactivity change of all quantified sites in the protein exceeded a threshold value (|log2Rsite – median(log2Rprotein)| > 0.6), the cysteine site was deemed to undergo a change in reactivity. If a given protein showed global, substantial changes in cysteine site reactivity (|median(log2Rprotein)| > 2), cysteine sites whose reactivity changed in the same direction as the protein (sign(log2Rsite) = sign(median(log2Rprotein))) were not deemed to undergo changes in reactivity. Finally, proteins for which the median change of cysteine-site reactivity was 2-fold vs. DMSO (|median(log2Rprotein)| > 1) were interpreted as changing in abundance. Note that, as discussed above, proteins were identified where changes in abundance coincided with site-specific changes in cysteine reactivity.

<u>Cysteine-site ligandability.</u> The ligandability of cysteine sites was determined independently under infected and uninfected conditions. For each condition, quantified PSMs were normalized using the corresponding control intensities as reference ([DMSO, no pathogen] = 100 or [DMSO, pathogen] = 100, respectively). Normalized intensities within each channel were then averaged across quantified PSMs associated to the same cysteine site to obtain raw site data. The relative blockade of IA-DTB labeling by the cysteine-directed scout fragment KB05 ^13^ was calculated as the difference between 100 and the average normalized PSM intensities for each cysteine site (e.g., an average normalized intensity of 20 corresponds to 80% IA-DTB blockade). In this analysis, lowest-variability quantified sites were those detected in 2 or more ABPP replicates and either (i) IA-DTB blockade ≥ 75% and CV ≤ 0.4 or (ii) IA-DTB blockade < 75% and sd ≤ 30. Channel values were averaged and grouped by protein. A cysteine site was deemed liganded by the scout fragment if IA-DTB blockade > 75% and at least one cysteine site in the same protein led to IA-DTB blockade < 25%.

### Luminescent Mtb infections and readings

THP-1 cells were differentiated in triplicate in a 96-well plate at 70,000 cells per well and infected at an MOI of three with Mtb-lux. Luminescence readings were taken at 0, 2, and 4 days after a PBS wash and media change to eliminate the contribution of extracellular bacteria.

### CFU assays

THP-1 macrophages were differentiated in 48-well TC-treated plates and infected at an MOI of one in triplicate. Cells were lysed at the indicated time-points with 0.5% Triton X-100 (Sigma) in water and serial dilutions were plated on 7H11 plates.

### Protein structures

All structures were downloaded as PDB files. Molecular graphics and analyses were performed with UCSF Chimera ^83^, developed by the Resource for Biocomputing, Visualization, and Informatics at the University of California, San Francisco, with support from NIH P41-GM103311.

## Statistical analysis

Significance of changes in cysteine reactivity was determined using the Kruskal-Wallis ANOVA (p ≤ 0.05) to compare MS signal between quantified cysteines within the same protein. Significance of changes in bacterial replication in THP-1 shRNA cell lines was determined using the Kruskal-Wallis ANOVA with multiple comparisons (p ≤ 0.05) to compare levels of bacterial replication in shRNA knockdown lines as compared to an infected WT THP-1 control. Significance of loss of MS signal dye to scout fragment binding was determined using an unpaired T-test with Welch’s correction

## Data availability

All mass spectrometry proteomics data have been deposited to the ProteomeXchange Consortium via the PRIDE partner repository ^84^ with the dataset identifier PXD067709.

## CRediT authorship contribution statement

John Neff: Conceptualization, Formal analysis, Investigation, Writing – original draft, Writing – review and editing. Kristen E. DeMeester: Conceptualization, Formal analysis, Investigation, Writing – original draft, Writing – review and editing. Paola Parraga: Investigation, Formal Analysis, Writing – review and editing. Radu Suciu: Investigation, Writing – review and editing. Melissa Dix: Investigation, Writing – review and editing. Gabriel Simon: Investigation, Writing – review and editing. Max A. Gianakopoulos: Investigation, Writing – review and editing. Bruno Melillo: Conceptualization, Formal analysis, Investigation, Writing – original draft, Writing – review and editing. Benjamin Cravatt: Conceptualization, Formal analysis, Funding acquisition, Project administration, Supervision, Writing – review and editing. Michael U. Shiloh: Conceptualization, Formal analysis, Funding acquisition, Project administration, Supervision, Writing – original draft, Writing – review and editing.

## Acknowledgements

This work was supported by the NIH (U19 AI142784) and the Jane Coffin Childs Memorial Fellowship (K.E.D). The authors thank members of the Shiloh and Cravatt labs for helpful suggestions and contributions.

## Abbreviations Used

ABPP: Activity Based Protein Profiling
CFU: colony forming unit
IA-DTB: iodoacetamide-desthiobiotin
PBMCs: human macrophages
MS: Mass Spectrometry
Mtb: *Mycobacterium tuberculosis*

---

**Supplementary ABPP Data Table 1.** All ABPP data (Tables S1-S18) are embedded within the supplementary Excel file. A table of contents is shared below.

| Supplementary Dataset |  |  |
| --- | --- | --- |
| Table | Tab Name | Description |
| S1 | experiments | List of analyzed ABPP experiments |
| S2 | workflow | Workflow for analysis of ABPP data |
| S3 | quantification | Number of cysteine sites and proteins quantified in ABPP experiments |
| S4 | all proteins | Summary of inferred changes in protein abundance, cysteine-site reactivity, and cysteine-site ligandability upon Mtb infection |
| S5 | abundance | Summary of inferred changes in protein abundance upon Mtb infection |
| S6 | reactivity | Summary of inferred changes in reactivity upon Mtb infection |
| S7 | THP (4 h) | List of inferred changes in protein abundance and cysteine-site reactivity upon Mtb infection (4 h post-infection, THP-1 cells) |
| S8 | THP (16 h) | List of inferred changes in protein abundance and cysteine-site reactivity upon Mtb infection (16 h post-infection, THP-1 cells) |
| S9 | PBMC (4 h) | List of inferred changes in protein abundance and cysteine-site reactivity upon Mtb infection (4 h post-infection, PBMCs) |
| S10 | PBMC (16 h) | List of inferred changes in protein abundance and cysteine-site reactivity upon Mtb infection (16 h post-infection, PBMCs) |
| S11 | REACTOME THP (16h) | REACTOME analysis results for proteins changing in abundance or site-specific cysteine reactivity in THP-1 cells at 16 h post-infection |
| S12 | REACTOME PBMC (16h) | REACTOME analysis results for proteins changing in abundance or site-specific cysteine reactivity in PBMC at 16 h post-infection |
| S13 | THP (4 h) ligandability | List of inferred changes in cysteine-site ligandability upon Mtb infection (4 h post-infection, THP-1 cells, KB05 scout fragment [200 μM, 1 h treatment]) |
| S14 | THP (16 h) ligandability | List of inferred changes in cysteine-site ligandability upon Mtb infection (16 h post-infection, THP-1 cells, KB05 scout fragment [200 μM, 1 h treatment]) |
| S15 | PBMC (4 h) ligandability | List of inferred changes in cysteine-site ligandability upon Mtb infection (4 h post-infection, PBMCs, KB05 scout fragment [200 μM, 1 h treatment]) |
| S16 | PBMC (16 h) ligandability | List of inferred changes in cysteine-site ligandability upon Mtb infection (16 h post-infection, PBMCs, KB05 scout fragment [200 μM, 1 h treatment]) |
| S17 | ligandability heatmap | Summary of inferred changes in cysteine-site ligandability upon Mtb infection |
| S18 | shRNA knock-down sequences | Gene names and oligo sequences for shRNA experiments |

